# T_1_-weighted fMRI in mouse visual cortex using an Iron Oxide Nanoparticle contrast agent and Ultrashort Echo Time (UTE) imaging at 9.4 T

**DOI:** 10.1101/2025.03.17.643844

**Authors:** Naman Jain, Saskia Bollmann, Kai-Hsiang Chuang, Jonathan R. Polimeni, Markus Barth

## Abstract

**Purpose:** This study aims to investigate the feasibility of using T_1_-weighted fMRI with an iron oxide nanoparticle contrast agent and Ultrashort Echo Time (UTE) imaging at 9.4T to measure functional hyperaemia in the mouse visual cortex. The goal is to capture positive signal changes in both the parenchyma and pial surface, and to test whether surface vessels respond during neuronal activation.

**Methods:** The study involved scanning of nine mice after administration of iron oxide-based superparamagnetic contrast agent (Molday ION) via the tail vein. Two functional imaging experiments were conducted: one to investigate the effect of echo time on the functional response, and another to characterize the impact of higher resolution on UTE functional contrast. Regions of interest (ROIs) were defined in the parenchyma and pial surface of the visual cortex.

**Results:** The administration of the contrast agent produced a bright-blood signal in the vasculature in structural MRI when using a UTE acquisition. Positive signal changes were observed at the shortest echo time (0.164 ms) in both the parenchyma (0.2% ± 0.08) and pial surface (0.2% ± 0.1 %), providing evidence that UTE fMRI experiments can detect changes in both pial and parenchymal vessels. Measurements using longer echo times (≥1 ms) showed negative signal changes. Higher spatial resolution resulted in increased percent signal change at the pial surface, suggesting less partial volume effects and better delineation of surface vessels.

**Conclusion:** The findings demonstrate that T_1_-weighted fMRI with UTE imaging and iron oxide nanoparticles captures positive signal changes across all vascular compartments, providing additional insights into the involvement of surface vessels during functional hyperemia.

## Introduction

The role different vascular compartments play during functional hyperaemia has been debated extensively in the field of functional neuroimaging (see, e.g., Drew, 2019; Kennerley et al., 2010; Mishra et al., 2016; Nippert et al., 2018; Rungta et al., 2018; Vaucher et al., 2000). *In vivo* measurements of various hemodynamic components can aid in addressing the question of how these compartments are engaged and interact during functional hyperemia, and thereby provide deeper insight into the vascular response to neuronal activity and improve the interpretation of functional MRI signals. Changes in cerebral blood volume (CBV) within arteries and small arterioles are of particular interest (Rungta et al., 2018; Schaeffer & Iadecola, 2021), as these constitute the ‘active’ component of the hemodynamic response, from which changes in cerebral blood flow and oxygenation follow (Boas et al., 2008; Buxton et al., 2004; Hillman et al., 2007; Kim et al., 2007; Lee et al., 2001; Stefanovic et al., 2008).

MRI-based CBV measurements in small-animal models typically use iron oxide-based contrast agents injected into the bloodstream (Berry et al., 1996; Hamberg et al., 1996; Kennan et al., 1998; Mandeville et al., 1998). The paramagnetic properties of iron oxide nanoparticles (Bjornerud et al., 2002; Kwon et al., 2015; Xiao et al., 2016) induce a dephasing of spins in and around blood vessels in gradient-echo-based acquisitions, creating a black-blood contrast and thus generate a signal decrease proportional to the increase in CBV (Mandeville & Marota, 1999) during functional hyperemia. CBV-fMRI has been shown to provide a higher contrast-to-noise ratio (Mandeville, 2012) and, because of the strong signal dephasing around large pial vessels, blood volume changes at the cortical surface are filtered out resulting in higher specificity to the parenchymal microvasculature (Mandeville & Marota, 1999). CBV-fMRI—with or without contrast agent—has been used to study brain function (Huber et al., 2021; Jin & Kim, 2006, 2008; Keilholz et al., 2006; Leite et al., 2002; Zhao et al., 2006) and showed improved spatial localisation (L. R. Huber et al., 2021; Jin & Kim, 2008, 2008; S. G. Kim et al., 2013; Mandeville & Marota, 1999) due to the apparent insensitivity to mid-sized and large vessels, especially the large draining veins on the pial surface of the cortex that dominate the BOLD contrast (Mandeville & Marota, 1999).

Optical imaging techniques such as two-photon microscopy also allow the observation of vascular responses to functional hyperaemia (Drew et al., 2011). Beyond changes in oxygenation, direct measurements of changes in vascular diameter of individual vessels—which is related to changes in vascular volume and thus CBV—can be made (Hillman et al., 2007). Two optical imaging studies (Drew et al., 2011; Tian et al., 2010) reported large diameter changes mostly in surface arteries, showing a discrepancy between CBV changes observed with optical imaging and those observed using iron-oxide nanoparticles as a contrast agent in fMRI. Note, however, that signal changes at the pial surface have been occasionally observed in CBV-fMRI studies (Donahue et al., 2006; Dresbach et al., 2023; L. R. Huber et al., 2021; Jin & Kim, 2008; Zhao et al., 2006).

In addition to their well-known T_2_* shortening effect, iron oxide nanoparticles also cause a marked T_1_-shortening (Girard et al., 2011; S. G. Kim et al., 2013). This can be exploited to create a bright-blood contrast when sufficiently reducing dephasing (T_2_*) effects by minimizing the echo time (TE), e.g., via zero echo time (ZTE) (MacKinnon et al., 2021) or ultra-short echo time (UTE) sequences (Gharagouzloo et al., 2015, 2017; Girard et al., 2011). Examples of structural vascular imaging were shown by Gharagouzloo and colleagues using a 3D UTE sequence. In a recent conference-proceeding article, one group has demonstrated the potential of combining an iron oxide-based superparamagnetic contrast agent and ZTE for fMRI showing positive signal changes in stimulated brain areas in rats (MacKinnon et al., 2021).

Motived by this, the goal of our study was to further investigate T_1_-weighted fMRI responses using an iron oxide-based superparamagnetic contrast agent and a short-TE acquisition. To achieve a high in-plane resolution while maintaining reasonably short TR, we chose a 2D UTE acquisition scheme. To compare our findings with those of previous studies (Girard et al., 2011; Han et al., 2015; Poplawsky & Kim, 2014; Rosen et al., 1991) and to better understand the transition from a T_1_-weighted image contrast at short echo times to a T_2_*-weighted image contrast at later echo times, we repeated the functional experiments across a range of echo times (from 0.164 ms to 10 ms). While in predominantly T_1_-weighted images blood vessels are expected to exhibit a bright signal, they are expected to appear dark in the predominantly T_2_*-weighted images as used in classical CBV imaging. Notably, CBV changes in large vessels should be preserved, since little-to-no dephasing occurs in the predominantly T_1_-weighted images. To better understand the contributions of different vascular compartments in each of these conditions, we analysed the task response in the parenchyma as well as the pial surface of the visual cortex, assuming that the parenchyma mainly contains capillaries and small arterioles and venules, and the pial surface predominantly larger vessels. Lastly, we also explored the possibility of even higher spatial resolution to reduce partial volume effects on the pial surface by performing the experiment at the shortest achievable echo time with an in-plane resolution of 0.1 mm.

## Material and Methods

### Animals and Ethics

All animal experiments were conducted with the approval of the Institutional Animal Ethics Committee at the University of Queensland. Imaging data from nine male C57BL/6J (age: 8– 10 weeks) mice were acquired.

For preparation, the animals were anaesthetized with 3% isoflurane mixed with air and oxygen in a 2:1 ratio until unconscious. The iron oxide-based superparamagnetic contrast agent Molday ION (BioPAL Inc., 5 mg Fe/ml) was administered into the tail vein by a senior animal technician with a dose of approximately 29.5 mg/kg. For sedation, a medetomidine bolus of 0.03–0.05 mg/kg was injected intraperitoneally.

For imaging, mice were positioned on an animal cradle, and their heads were fixed using a bite bar and ear bars to limit movement. Vital parameters such as respiration rate, body temperature, heart rate and blood oxygenation levels were continuously monitored by a veterinary support officer throughout the imaging session. A respiration sensor was placed beneath the belly of the mice, a rectal temperature probe was used to monitor body temperature, and a tail-cuff SpO_2_ sensor was employed to measure both heart rate and blood oxygenation level. Body temperature was maintained at 36–37°C by using a warm-water cushion whose temperature could be adjusted based on the body temperature of the mice. Respiration rate was maintained at 120–150 breaths per minute, heart rate at 210–230 beats per minute and SpO_2_ levels at >96% by adjusting the isoflurane concentration between 0.2 % and 0.5 %. Medetomidine was supplied constantly during imaging through an intraperitoneal infusion at 0.085–0.1 mg/kg/hour.

### MRI Data Acquisition

Imaging data were acquired on a Bruker Biospec 9.4T preclinical MRI system (Ettlingen, Germany) controlled by a 30 USR Avance III console running Paravision 6.0.1. Maximum gradient strength and slew rate were 660 mT/m and 6000 mT/m/s, respectively. An 86-mm quadrature volume coil was utilised for transmission, and an in-house developed 10-mm ellipsoid surface coil was used for reception.

An anatomical reference image was acquired using a T_2_-weighted Turbo-RARE sequence with a TE of 55 ms, a voxel size of 0.1×0.1×0.25 mm^3^, 24 slices, and three averages with a total acquisition time of 6 minutes and 36 seconds. A map of the vasculature was obtained in five mice using a 3D UTE sequence with a TE of 13 μs, a repetition time (TR) of 5 ms, an isotropic nominal resolution of 0.1 mm, 115,432 radial spokes, a final matrix size of 192×192×192 and a total acquisition time of 9 minutes and 37 seconds.

Two different functional imaging experiments were performed, and animals were divided into two groups with different spatial resolutions: *Group 1 – High-Resolution* used 0.2 × 0.2 × 0.5 mm^3^; and *Group 2 – Ultra-High-Resolution* used 0.1 × 0.1 × 0.5 mm^3^. In Experiment 1, the effect of echo time on the functional response was investigated in the five mice of Group 1 by acquiring functional data with a voxel size of 0.2 × 0.2 × 0.5 mm^3^ at five different echo times using either a 2D UTE (0.164 ms, 0.5 ms, 1 ms) or a 2D FLASH (4 ms and 10 ms) sequence (see Table 1). To minimize T_2_* contamination in the UTE acquisitions, the shortest possible readout durations of 0.32 ms for the high-resolution and 0.96 ms for ultra-high-resolution acquisition were used.

**Table 1:** Comparison of protocol parameter values between UTE and FLASH acquisitions.

|  | **2D UTE**<br>**(ultra-high-**<br>**resolution)** | **2D UTE**<br>**(high-resolution)** | **2D FLASH** |
| --- | --- | --- | --- |
| **TE** | 0.164 ms | 0.164 ms, 0.5 ms<br>and 1 ms | 4 ms and 10 ms |
| **TR** | 4.968 ms | 4.968 ms | 55.556 ms |
| **VOLUME TR** | 3 s | 3 s | 3 s |
| **NUMBER OF SLICES** | 1 | 2 | 8 (TE=4 ms),<br>4 (TE=10 ms) |
| **RESOLUTION** | $0.1 \times 0.1 \times 0.5 \text{ mm}^3$ | $0.2 \times 0.2 \times 0.5 \text{ mm}^3$ | $0.2 \times 0.2 \times 0.5 \text{ mm}^3$ |
| **SLICE GAP** |  | 0.1 mm | 0.1 mm |
| **MATRIX SIZE** | $192 \times 192$ | $96 \times 96$ | $64 \times 54$ |
| **NUMBER OF RADIAL**<br>**SPOKES** | 604 | 302 | -NA- |
| **READOUT**<br>**BANDWIDTH** | 100,000 Hz | 100,000 Hz | 50,000 Hz |
| **READOUT DURATION** | 0.96 ms | 0.32 ms | 1.28 ms |

To ensure a sufficiently short volume-TR of 3 s compatible with the timing of the hemodynamic response, the coverage of the 2D UTE acquisition was limited to two slices, and the number of slices in the FLASH acquisitions was set to either eight (TE = 4 ms) or four (TE = 10 ms) to match the volume TR. A system-calculated RF excitation pulse with a flip angle (FA) of 15° was used to acquire the FLASH images, and to achieve shortest TE for the UTE images a Half-Gauss RF excitation pulse with an FA of 20° was used with the flip angle optimisation based on previous work (Gharagouzloo et al., 2015). The FLASH acquisition with TE = 4 ms was performed first to serve as a functional localiser, and the two slices with the strongest responses were chosen to plan all subsequent functional scans. The order of the remaining scans was pseudo-randomly alternated for each mouse. Before each 2D UTE acquisition, an online k-space trajectory adjustment with 32 averages was performed. As a control, one additional mouse was scanned without contrast injection.

In Experiment 2, the effect of higher spatial resolution was investigated in the four animals of Group 2. First, the FLASH sequence with 4-ms TE was used as a functional localiser. Then, single-slice 2D UTE images with the shortest TE of 0.164 ms and a voxel size of 0.1 × 0.1 × 0.5 mm^3^ were acquired. To compensate for the reduction in SNR due to the smaller voxel volume, we acquired more runs to provide noise cancellation through data averaging (two runs in two mice, and four runs in two mice).

### Stimulation Paradigm

The visual stimulus consisted of a blue flashing light delivered through an optical fibre (frequency 5 Hz, pulse duration 10 ms, power <1 mW). To minimise the risk of eye damage, the optical fibre was pointed towards the rear of the scanner bore to avoid projecting light directly into the eyes of the animals. The stimulation paradigm utilised a block design, with eight blocks of 30-second rest interspersed with seven blocks of 30-second stimulation. In total, each run lasted 7 minutes and 30 seconds, and a 5-minute rest period was inserted between runs. In Experiment 1, one run was acquired per TE and animal, whereas in Experiment 2, one run for the functional localizer and between two to four runs per animal were acquired for the shortest TE (0.164 ms).

### Activation Map Estimation

FSL (Jenkinson et al., 2012; Smith et al., 2004) (https://fsl.fmrib.ox.ac.uk/fsl) and AFNI (Cox, 1996; Cox & Hyde, 1997) (https://afni.nimh.nih.gov/) software packages were utilised for data processing. The first three volumes of each run were discarded to ensure the MR signal had reached steady state. Motion correction was applied using *3dvolreg* from the AFNI package, and Fourier interpolation was used for resampling. The middle volume was selected as the reference image for motion correction. To enhance the signal-to-noise ratio (SNR) for extracting activation maps, spatial smoothing with a full-width-at-half-maximum equivalent to two voxels was applied, i.e., 0.4 mm for the high-resolution data and 0.2 mm for the ultra-high-resolution data. We utilized a hemodynamic response function (HRF) tailored to the mouse cortex; the HRF was modelled as a Gamma-variate function with phase 0 s, standard deviation 2 s, and mean lag 4 s. Temporal derivatives were added to the model, and a high-pass filter with a cut-off of 90 s was applied. Data were analysed using a General Linear Model, as implemented in the GLM Setup package in FSL.

### ROI Definition and Analysis

Percent signal changes of the fMRI data were investigated in two regions of interest (ROIs), namely the parenchyma and the pial surface of visual cortex. The ROIs were defined on the TurboRARE anatomical reference image using the Paxinos and Franklin mouse brain atlas (Paxinos & Franklin, 2001). The parenchymal ROI was placed centrally in cortical grey matter, leaving a two-voxel-wide gap between the ROI and the cortical surface. A one-voxel-thick pial surface ROI was then placed above the parenchymal ROI. On average, 21 voxels (in native fMRI space) contributed to the parenchymal ROI and 9 voxels (in native fMRI space) contributed to the pial surface ROI.

To minimise the need for manual realignment, similar image contrasts within each sequence type, UTE (TE: 0.164 ms, 0.5 ms and 1 ms) or FLASH (TE: 4 ms and 10 ms), were first concatenated and then motion corrected (as shown in Figure 1a). This resulted in one mean functional image per sequence type, which was then manually aligned to the anatomical reference image using ITK-Snap (Yushkevich et al., 2006) producing a rigid transformation matrix. The AFNI *3dresample* function was then used to resample the functional time series into the space of the anatomical reference image. To avoid bias when aligning the mean functional and structural images due to the signal decay at the surface of the images with longer TE, the skin and fat signal of the skull were used as landmarks.

**Figure 1:**
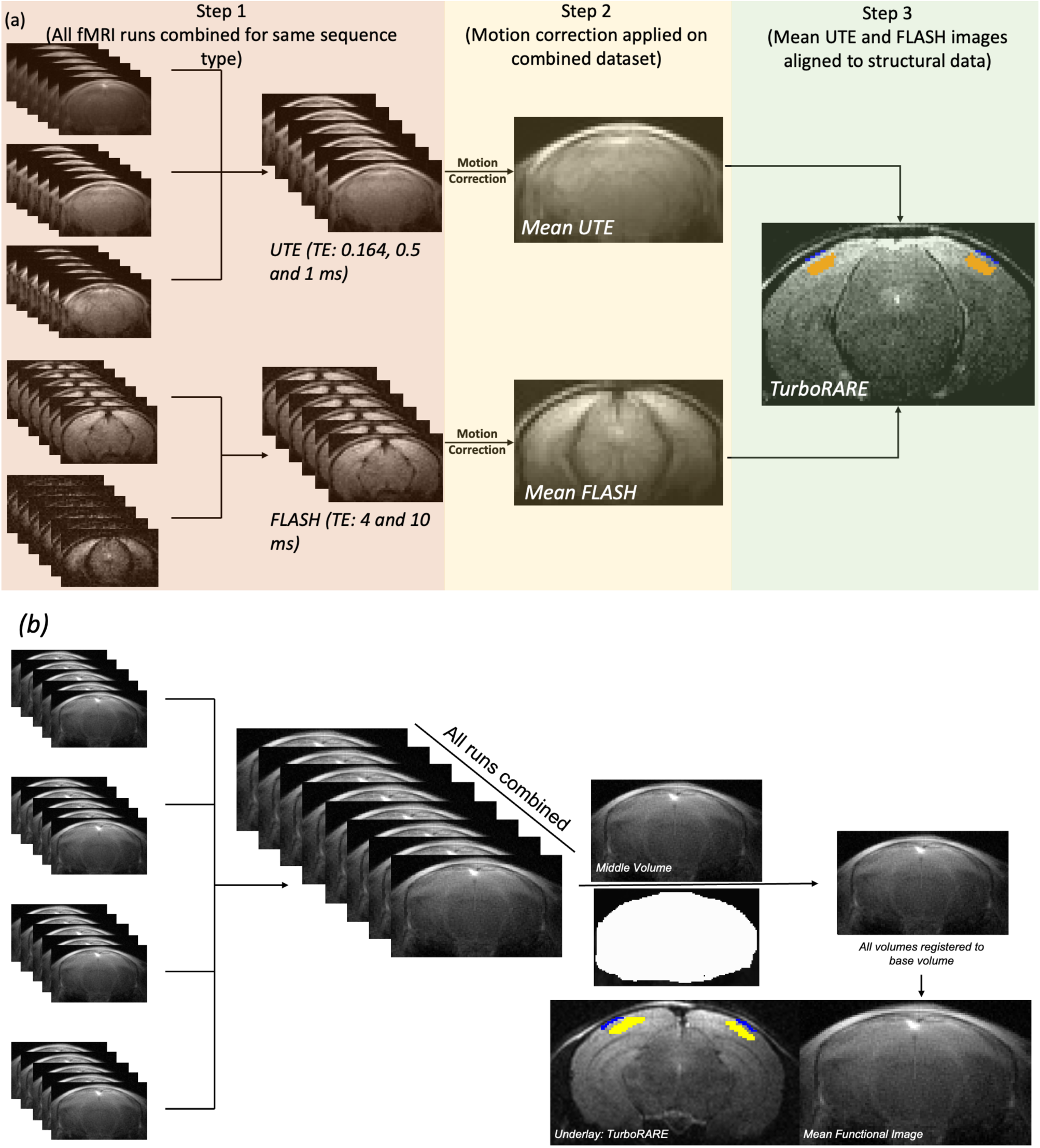
Data Analysis Pipeline for (a) High resolution data and (b) Ultra-high resolution data. For the high-resolution data in Step 1, all functional data from the same sequence type, i.e. either UTE or FLASH, were concatenated and then motion corrected in Step 2. In Step 3, the mean image of the combined data set was then manually aligned to the structural image on which the ROIs were defined. For the ultra-high resolution data, all runs of the UTE images acquired at 0.164 ms were concatenated and then motion corrected using a brain mask to improve the motion parameter estimation. The resulting mean image was then manually aligned to the reference structural image.

To estimate the trial responses within the ROIs for each individual TE value, the demeaned time series across the 7 trials in each run and within each ROI were averaged. Percent signal change was then computed by subtracting and then dividing the averaged signal time course by the baseline signal level (approximated as the mean intensity of the last 6 data points immediately before stimulus onset in all trials for each run). From this, the average percent signal change was computed by averaging across 35 time points acquired during the stimulus presentation.

For comparison, we also computed the absolute image intensity in the pial surface ROI relative to the image intensity within the parenchyma of the visual cortex by scaling the average image intensity values in the parenchymal ROI to ‘1’ and then applying the same scaling factor to the pial surface ROI. This enabled a comparison of differences in percent signal change relative to differences in image intensity.

### Analysis of the ultra-high-resolution UTE data

The analysis of the UTE data acquired with 0.1-mm in-plane resolution followed a similar approach as outlined above. Since only a single 2D image slice was acquired, *3dAllineate* was used for motion correction as it supports alignment of single 2D images. The same procedure as for the high-resolution data was employed to compute the trial responses for both ROIs (Figure 1b).

## Results

The administration of the iron oxide-based superparamagnetic contrast agent reduced the T_1_ of blood and produced a bright-blood signal in the vasculature when using the UTE acquisition, as previously shown (Gharagouzloo et al., 2015). Figure 2 illustrates this effect, showing a whole-brain maximum-intensity projection of a 3D UTE image, acquired with a TE of 13 μs, from one mouse.

**Figure 2:**
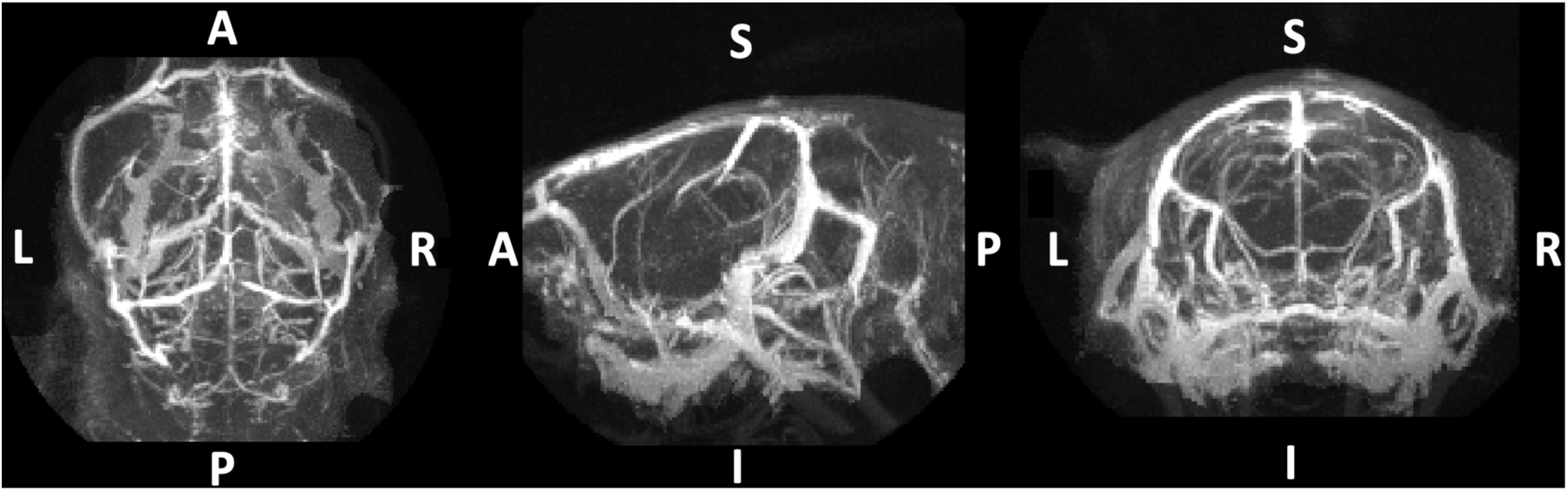
Example of a Maximum-Intensity Projection of a whole mouse brain acquired using a 3D UTE sequence at 9.4 T after iron oxide-based superparamagnetic contrast agent injection, showing high signal intensities in the vasculature.

When comparing the functional responses to visual stimulation across five different echoes times ranging from 0.164 ms to 10 ms, positive signal changes were found in all mice at the shortest echo time, and negative signal changes were found at longer echo times (TE ≥ 1 ms) (Figure 3). The data acquired with the two longest echo times exhibit the expected negative signal change with activation (Keilholz et al., 2006; Kim et al., 2013; Mandeville & Marota, 1999) in mouse visual cortex, whereas the activation region observed in the data acquired at TE = 1 ms included visual cortex but also the superior sagittal sinus. No clear activation pattern was seen in data acquired at the intermediate TE of 0.5 ms. As a control, we also repeated the experiment without contrast agent and acquired data using the same acquisition protocols. There, positive BOLD activation patterns in visual cortex were present for the two longest TE values, but no consistent responses were found at shorter echo times (TE ≤1 ms).

**Figure 3:**
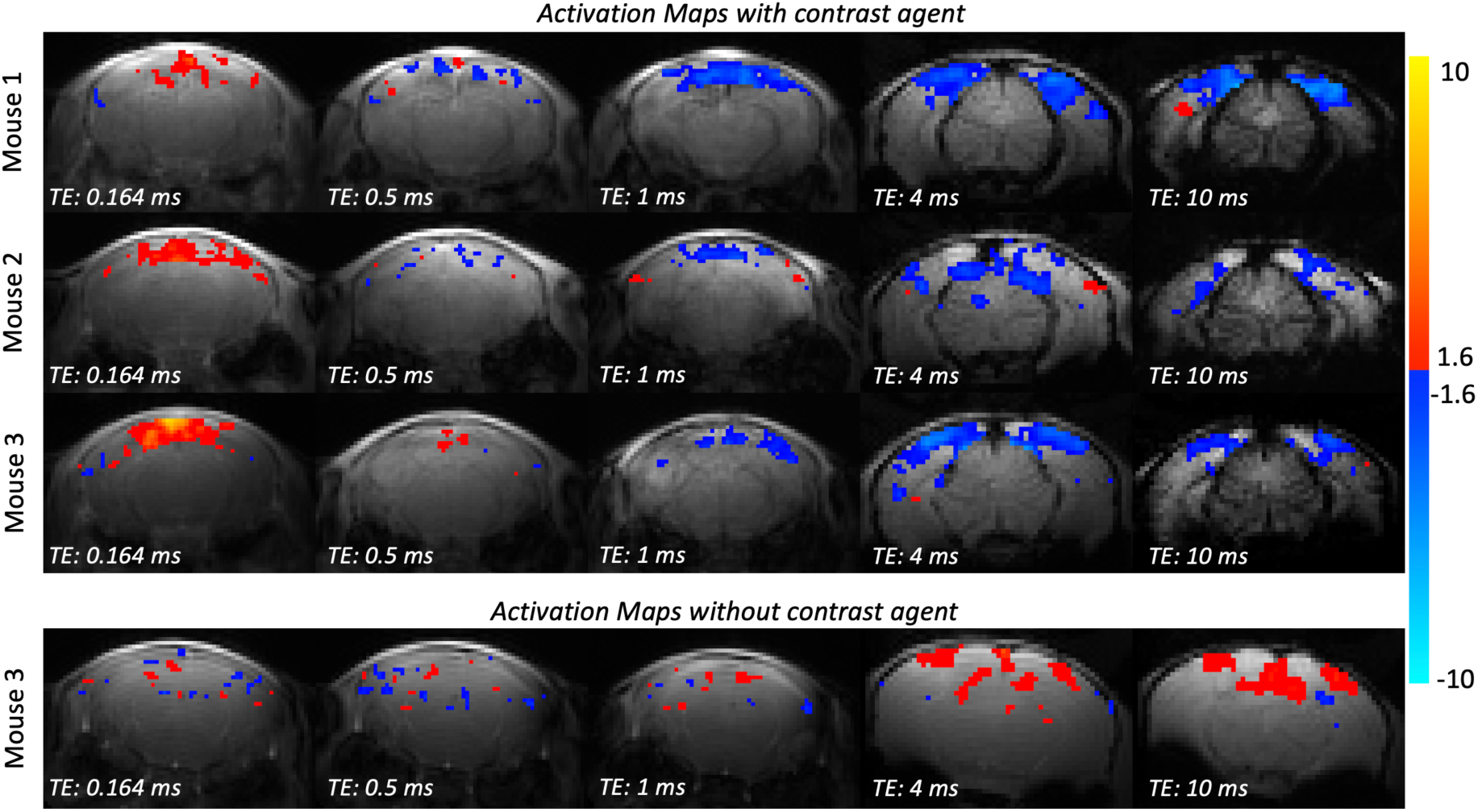
Activation maps (p < 0.05) from three animals in Group 1 showing responses to visual stimulation across five different TEs acquired with a 2D UTE sequence (0.164 ms, 0.5 ms, and 1 ms TE) or a 2D FLASH sequence (4 ms and 10 ms TE) after contrast agent administration (rows 1–3). The data acquired with the shortest TE exhibit positive signal changes (positive z-values overlayed in hot colorscale), while data using longer TEs exhibit negative signal changes (negative z-values overlayed in cold colorscale). The bottommost row shows activation maps (p < 0.05) for the control experiment without contrast agent. For data acquired with short echo times (≤1 ms), no distinct activation was found, whereas in data acquired at longer echo times, positive signal changes in visual cortex were observed consistent with a BOLD response.

To investigate the effect of echo time on the response strength within the parenchyma of the visual cortex and at its surface, percent signal change across the five TE values for each ROI were compared (Figure 4a). In the data acquired with the shortest TE, small positive signal changes of similar magnitude (mean: 0.26%, averaged across the 5 mice of Group 1) in both ROIs were observed. Within both the pial surface and parenchyma ROIs, the percent signal change consistently decreased with TE and reached a negative value of about −1.7% at the longest echo time. Figure 4b shows a steady decrease in image intensity of the pial surface ROI relative to the parenchymal ROI with increasing TE values.

**Figure 4:**
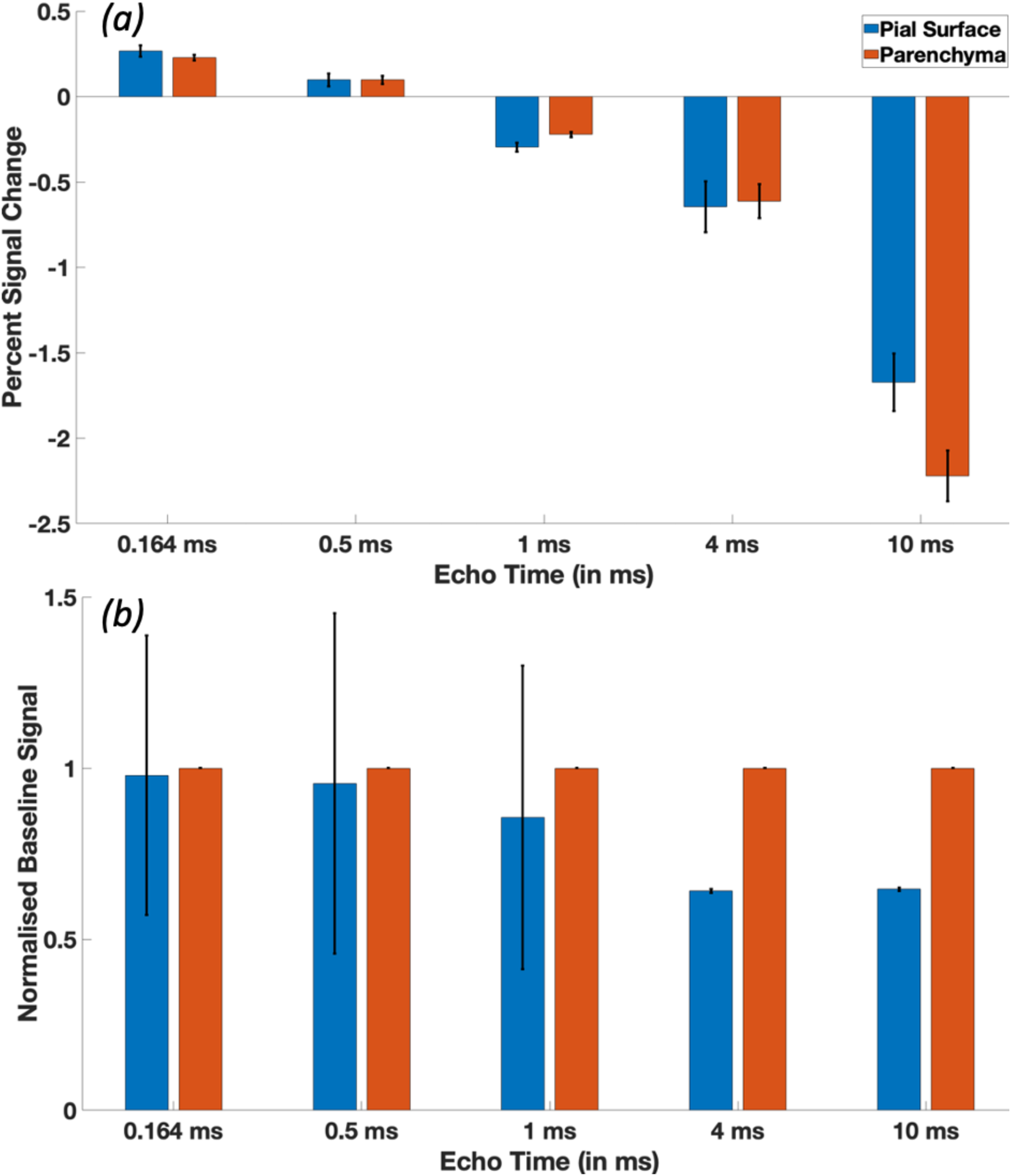
(a)Percent signal change (n = 5) as a function of TE after contrast injection measured with a 2D UTE (TE ≤ 1 ms) or 2D FLASH (TE > 1 ms) sequence. At short TE values, positive percent signal changes were small, but of similar magnitude on the pial surface (0.26% ± 0.03%) and in the parenchyma (0.22% ± 0.01%). With increasing TE, percent signal change decreased and reached strong negative values of similar magnitude (pial surface: −1.67% ± 0.16% and parenchyma: −2.22% ± 0.14%) at the longer echo times (TE >1 ms) in both the parenchyma and on the surface. Error bars indicate standard error of the mean across five subjects. (b) Normalized image intensities averaged within both ROIs, scaled such that values in the parenchymal ROI always equal ‘1’, show a steady decrease in the pial surface ROI with TE value.

The trial-averaged fMRI responses are presented in Figure 5 both for the predominantly T_1_-weighted data (Figure 5a) corresponding to the UTE acquisition with the shortest TE value and for the predominantly T_2_*-weighted data (Figure 5b) corresponding to the FLASH acquisition with the longest TE value. Responses are plotted for both the parenchyma and pial surface ROIs. As seen in Figure 4, these responses show that a clear positive signal change is seen in the T_1_-weighted data and clear negative signal change is seen in the T_2_*-weighted data. Both the amplitudes and timings of the responses seen in the parenchymal and pial surface ROIs are comparable within each acquisition, although subtle differences in timing may be appreciated between the responses measured with the T_1_-weighted data and the T_2_*-weighted data.

**Figure 5:**
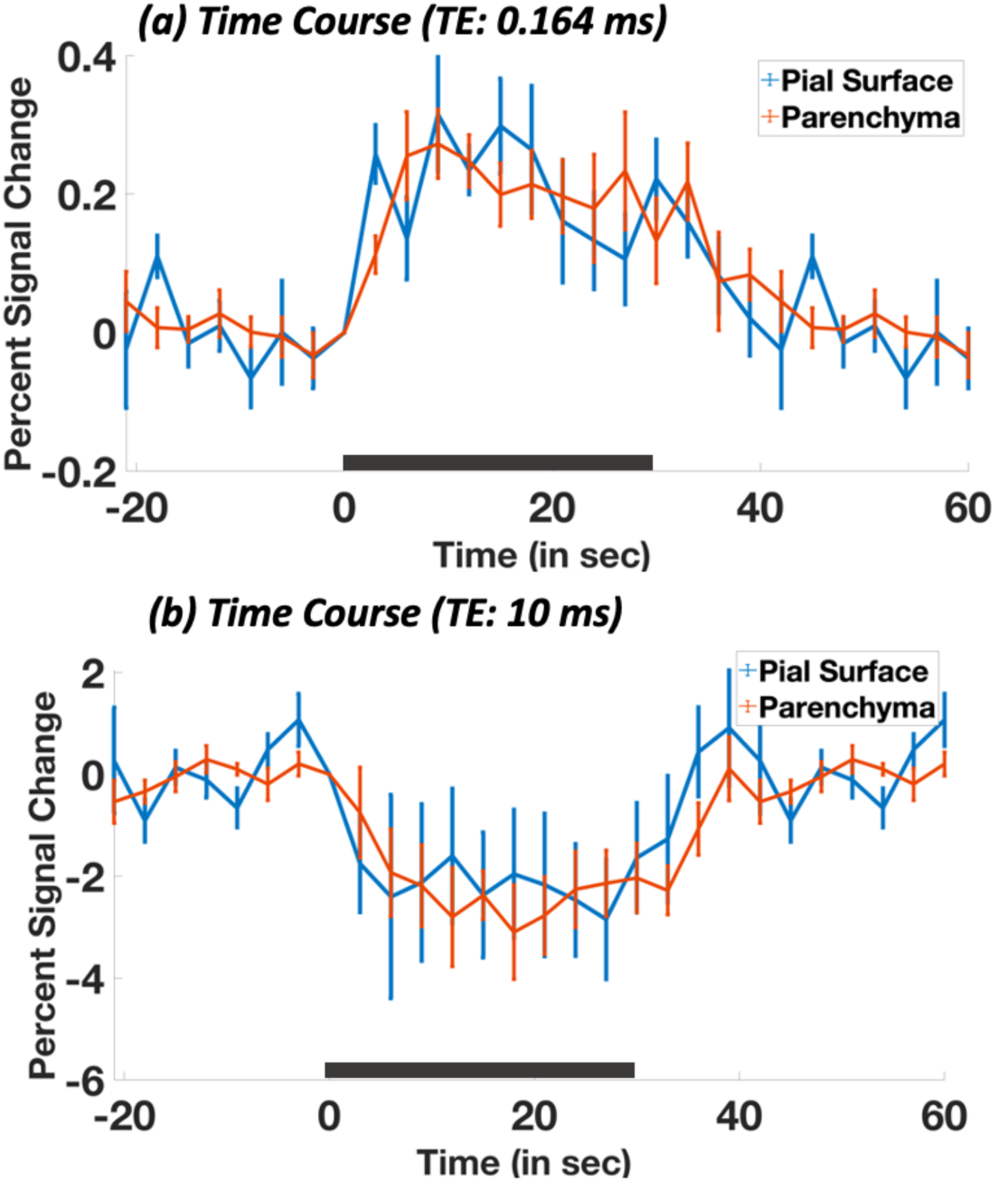
T Average time courses (n=5) within the parenchymal (orange) and pial surface ROI (blue) estimated from data acquired at the shortest (a) and longest (b) echo times. Error bars indicate standard error of mean across five subjects. Black bar indicates stimulus ON duration.

Figure 6a shows an example activation map of a UTE acquisition at ultra-high-resolution (0.1-mm) depicting significant functional activation throughout the cortex (Figure 6a); for reference, Figure 6b shows the two ROIs overlaid on ultra-high-resolution UTE-fMRI data. Figure 6c and d show the trial-responses for both the high- and ultra-high-resolution fMRI data both in the parenchyma and pial surface ROIs. In data averaged within the parenchymal ROI, we observed a similar response amplitude and shape for both imaging resolutions (Figure 6d), whereas in data averaged within the pial surface ROI the response amplitude of the ultra-high-resolution acquisitions (0.1 mm) was twice as high as the response compared to the high-resolution acquisitions (0.2 mm), but also highly variable across mice (see Discussion).

**Figure 6:**
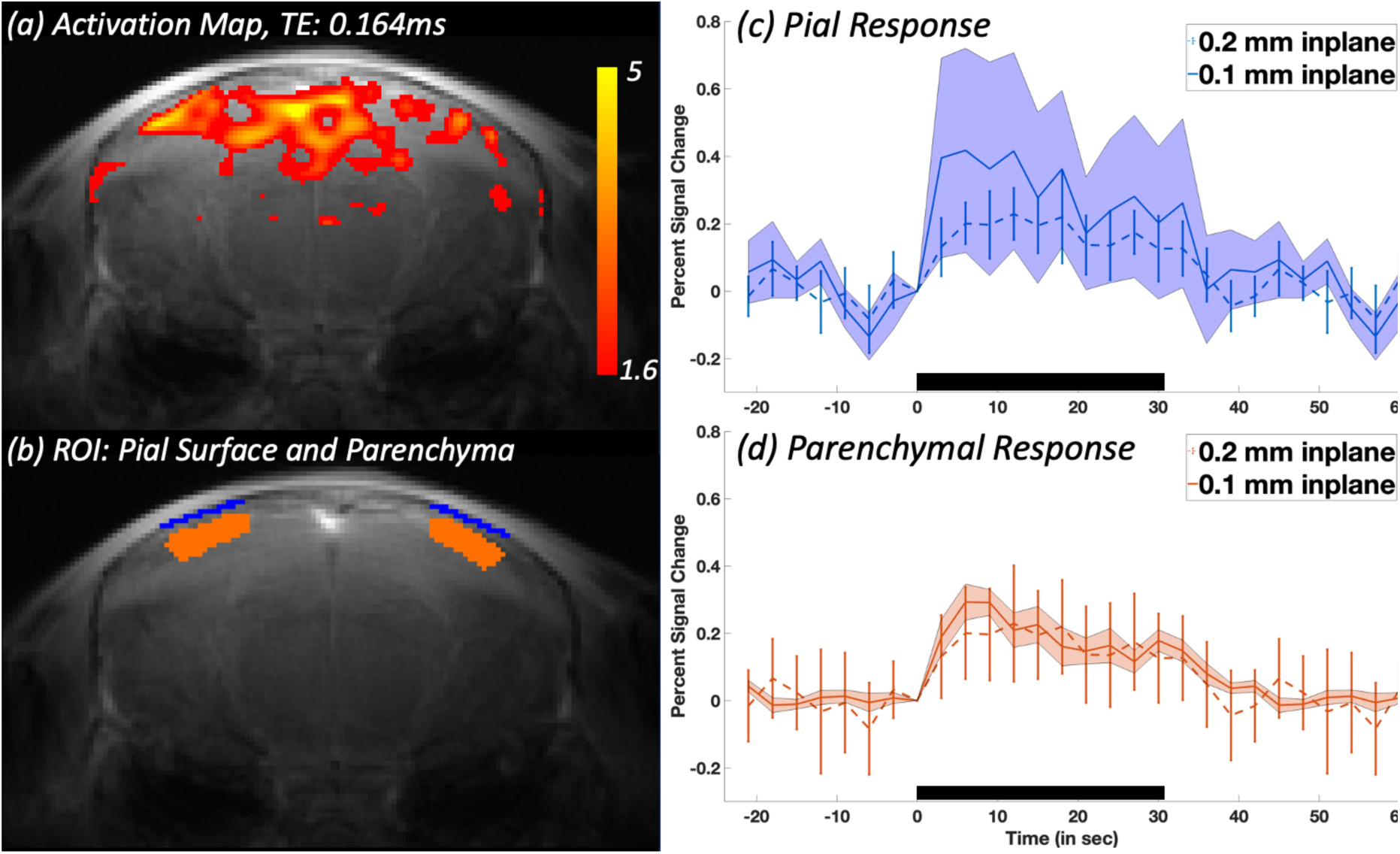
Comparison of UTE fMRI responses between the high-resolution and ultra-high-resolution fMRI acquisitions. (a) Example activation maps across one subject at ultra-high resolution. (b) Regions of Interest at Pial Surface (blue) and Parenchyma (orange). (c and d) Comparison of average time courses extracted from the pial surface (c) and parenchyma (d) ROIs using UTE, TE = 0.164 ms between both high- and ultra high-resolution fMRI data (Groups 1 and 2). The trial-responses are plotted as percent signal change averaged across all subjects (n=5 in Group 1, n=4 in Group 2) for the pial surface (Mean PSC: 0.47% ± 0.11 %) and parenchyma (Mean PSC: 0.21% ± 0.02 %) ROIs across both spatial resolutions: 0.2-mm resolution (dashed line) and 0.1-mm resolution (solid line). Black bar indicates stimulus duration; error bars represent standard error of the mean across subjects.

## Discussion

In this study we investigated the characteristics of fMRI signal changes after injection of the iron oxide-based superparamagnetic contrast agent Molday ION using a UTE acquisition scheme. This combination of MION and a UTE acquisition has been previously used in vascular anatomical imaging (Gharagouzloo et al., 2015) where it was shown to generate a bright-blood signal due to the T_1_-shortening property of MION when injected into the blood stream (as also shown in Figure 2). This is in distinction to “classical” iron oxide nanoparticle experiments (Girard et al., 2011; Han et al., 2015; Keilholz et al., 2006; S. G. Kim et al., 2013; Mandeville, 2012; Qiu et al., 2012) that use the T_2_*-shortening properties of the contrast agent in combination with T_2_*-weighted MRI acquisitions.

### Effect of Echo Times on Contrast Mechanisms

When varying the TE from 0.164 ms to 10 ms, we could observe the expected reversal in blood-tissue contrast from bright blood, i.e., T_1_-weighted, to dark blood, i.e., T_2_*-weighted, at around 1 ms (Figure 3). We also observed the corresponding reversal in the sign of the signal change with activation. The functional response in the T_2_*-weighted images was fairly well localised to the visual cortex (4 ms and 10 ms in Figure 3), whereas the response in the T_1_-weighted image (0.164 ms in Figure 3) was more spread out and also less consistent across mice. Further, the magnitude of the T_2_*-weighted percent signal changes were much higher than the T_1_-weighted signal change (Figure 4), indicating higher sensitivity and potentially lower noise variance, partly explaining the higher consistency. Note that we employed a dedicated processing pipeline (Figure 1) to minimise variance due to ROI selection by marking only one parenchymal and one pial ROI per mouse, aligning each sequence type (UTE or FLASH) across TEs and then having to align only two (instead of five) mean UTE or FLASH images manually to the structural reference image. Due to the limited coverage (two or one slice(s), respectively) and the vastly different image contrasts (T_1_ to T_2_ or T_2_* to T_2_), this step could only be successfully performed manually. Therein, we paid particular attention to features outside of the brain like fat and skull signal to mitigate the artefactual shift in the location of the pial surface due to the signal decay in the FLASH images.

While we could show the expected reduction in image intensity on the cortical surface (Figure 4b) presumably due to the dephasing effects around large vessels, our comparison of the percent signal changes at the pial surface and within the parenchyma did not show higher percent signal changes in the parenchyma (Figure 4a). One potential explanation for this observation is that computing relative signal change (i.e., ΔS/S) tends to amplify responses at the pial surface where the baseline signal (S in the denominator) is typically lower with increasing echo times. Computing relative signal change, however, allows to compare across different echo times.

The temporal characteristics of the parenchymal and pial response at the longest and shortest echo time appeared very similar (Figure 5); however the percent signal change in the T_1_-weighted fMRI data was much lower compared to the T_2_*-weighted images. While there is no comparable study for the T_1_-weighted data, at the longer echo time (TE: 10 ms) Zhao and colleagues observed a similar percent signal change on the pial surface and within the parenchyma in cats (2006).

Interestingly, numerous studies have reported a rapid and transient increase in arterial diameter in response to sensory stimulation, which may contribute to the observed response at the pial surface (Hua et al., 2019; Kennerley, Mayhew, et al., 2010; T. Kim & Kim, 2011, 2011; Lee et al., 2001; Zong et al., 2012). Notably, Drew et al. (2011) also documented changes in venous diameter with longer duration stimuli in vessels near the pial surface, which is particularly relevant given that our signal reflects total cerebral blood volume (CBV), suggesting contributions from both dilating arteries and veins at the pial surface.

### Effect of Spatial Resolution on Signal Changes

When increasing the in-plane resolution to 0.1-mm (Figure 6), we found similar activation patterns as see in the 0.2-mm data (Figure 3). The estimated percent signal change within the parenchyma was also comparable between both resolutions. However, the percent signal change at the pial surface increased when reducing the voxel size. The observable difference in the response on the pial surface is consistent with a reduced partial volume effect. However, to fully resolve individual pial arteries, voxel sizes of 34 µm to 41 µm in mice (Sekiguchi et al., 2014) and 50 µm to 300 µm in humans would be necessary. While initial studies have shown the potential to resolve these anatomically (Bollmann et al., 2022), adapting this anatomical acquisition into a functional acquisition would be needed; thus, resolving these small vessels both spatially and temporally remains challenging. Accordingly, CBV-fMRI may continue to rely on partial-volumes models for interpretation and quantification (Donahue et al., 2006). Importantly, while a two-compartment model describing grey matter and blood volume changes may be sufficient within the parenchyma, volume changes in cerebrospinal fluid and the potential dilation of grey matter with activation also need to be considered on the surface (Krieger et al., 2012).

### Contribution of blood flow

In previous studies, a large 3D volume was excited to limit signal enhancements flowing into the imaging volume of interest (Dechent et al., 2011), whereas here we only used one or two thin slice(s). Thus, the observed signal changes constitute a combination of both CBV changes and inflow effects. In our T_1_-weighted data, activation would be expected to increase CBV and inflow velocity, and so both physiological effects would contribute to an increase in signal and thus a positive response to activation. Indeed, at the shortest echo time we observed a positive response also in the superior sagittal sinus. Future studies could address this confound by either increasing the coverage or applying additional saturation pulses to the non-imaged regions to reduce inflow and isolate the CBV component.

### Involvement of Different Vascular Compartments

Commonly, the decrease in signal with activation is interpreted as being proportional to increases in CBV (S. G. Kim et al., 2013). In our study when employing longer TE (> 1ms) we were able to replicate this behaviour. However, when employing an ultra-short echo time using a 2D UTE sequence, we instead observed positive functional signal changes upon visual stimulation. One possible interpretation of this positive signal change is also as an increase in CBV, as suggested in the study of Gharagouzloo and colleagues (Gharagouzloo et al., 2017). One advantage of mapping changes in CBV using T_1_-weighting is the absence of the ‘vascular filter’ present when using T_2_*-weighting that removes contributions from large vessels via strong dephasing at the cortical surface attributable to strong susceptibility effects (T. Kim & Kim, 2011). This filter present in conventional T_2_*-weighted contrast-enhanced CBV-fMRI may be beneficial for studies targeting small vessels in the parenchyma (L. Huber et al., 2015; Jin & Kim, 2008; S. G. Kim et al., 2013; Rivera-Rivera et al., 2019) to maximise neural specificity, but may also obscure the involvement of larger macrovessels in functional hyperemia if the aim is to understand the neuro*vascular* response. Conversely, our approach based on T_1_-weighting is sensitive to all vessels, and thus reflects total CBV and not only microvascular CBV, and can provide a more complete picture of the vascular response.

## Conclusions

In this study, we investigated the feasibility of conducting fMRI experiments using a T_1_-weighted contrast combing UTE imaging and iron oxide nanoparticles in mice. Our findings demonstrate that, when echo times are sufficiently short as to minimize T_2_* weighting, a bright-blood signal following the administration of a contrast agent is observed, as well as a positive fMRI signal change with activation. At longer echo times, we observed a negative signal change across all regions of interest (ROIs), as shown in previous studies. The positive fMRI signal changes seen in the parenchyma and on the pial surface suggest an involvement of surface vessels during functional hyperaemia. These positive functional signal changes are consistent with blood volume responses to stimulation, however further studies are needed to disentangle possible contributions from inflow effects.

## Acknowledgements

This work was supported by an Australian Government Research Training Program Scholarship, NHMRC-NIH BRAIN Initiative Collaborative Research Grant APP1117020 and the NIH NIMH *BRAIN Initiative* grant R01-MH111419. Further, the National Imaging Facility (NIF) at the Centre for Advanced Imaging is acknowledged. I thank Dr. Nyoman Kurniawan for help with MRI scanner operation, Dr. Heidi-Niland Rowe and Barb Arnts from UQBR for support with data collection, and Shreya Jain help with analysis software development.

